# VolcanoR - web service to produce volcano plots and do basic enrichment analysis

**DOI:** 10.1101/165100

**Authors:** Vladimir Naumov, Ivan Balashov, Vadim Lagutin, Pavel Borovikov, Alexey Alexeev

## Abstract

Summary: We introduce VolcanoR - web based tool to analyse results of differential gene expression. It takes a table containing gene name p-value and foldChange as input data. It can produce publication quality volcano plots, apply different p-value and fold change thresholds and do basic GeneOntology and KEGG enrichment analysis with selected gene set. For now it supports H.sapiens, R.norvegicus and M.musclus.

Availability and Implementation: VolcanoR is wtitten using R Shiny framework. It is publically available at http://volcanor.bioinf.su or stand-alone application, that can be downloaded at https://github.com/vovalive/volcanoR

## Text

### Introduction

Differential gene expression analysis is widespread in modern bioscience. It is based on both RNA-seq or microarray gene expression analysis. Groups comparsion results are p-value and fold change for each gene analysed. Such results can be produced by multiple proprietary software products like Illumina GenomeStudio or Affymetrix Expression Console. It can also be produced right from NCBI GEO using easy-to use GEO2R software (Barrett et al., 2013). Volcano plot is one of quality control plots, that is used after differential expression analysis is performed. It shows relation between fold change and statistical confidence. Volcano plot is a clear and simple way to assess the results of the analysis. In VolcanoR it’s easy to apply different p-value and fold change thresholds and get nice visualization and gene set overrepresentation analysis using GeneOntology(Gene Ontology Consortium, 2015) and KEGG(Kanehisa, 2000) databases. Now we have versions for H.sapiens, R.norvegicus and M.musculus

### Implementation

VolcanoR is written in R shiny(Chang *et al.*, 2017) framework and several bioconductor libraries for data manipulation (Wickham and Francois, 2016), plotting (Wickham, 2009; Slowikowski, 2016; Warnes *et al.*, 2016), annotation (Carlson, 2016a, 2016c, 2016b) and gene set enrichment (Yu et al. 2012).

Interface contains two panels - left one for data uploading and parameters adjustment and right for results.

The process of interaction is quite easy. First it’s needed to choose an organism of interest from a dropdown menu. Then user uploads differential expression data in tab separated format, we also have several examples. After data uploading it’s possible to adjust p-value and fold change thresholds and get volcano plot with significant genes labeled. It’s also possible to do GO and KEGG Enrichment Analysis of a gene set. The resulting Volcano plot, table with enrichment results and significant genes will appear on the right panel.

The tool is publicly available at http://volcanor.bioinf.su and in GitHub repository https://github.com/vovalive/volcanoR1.

---

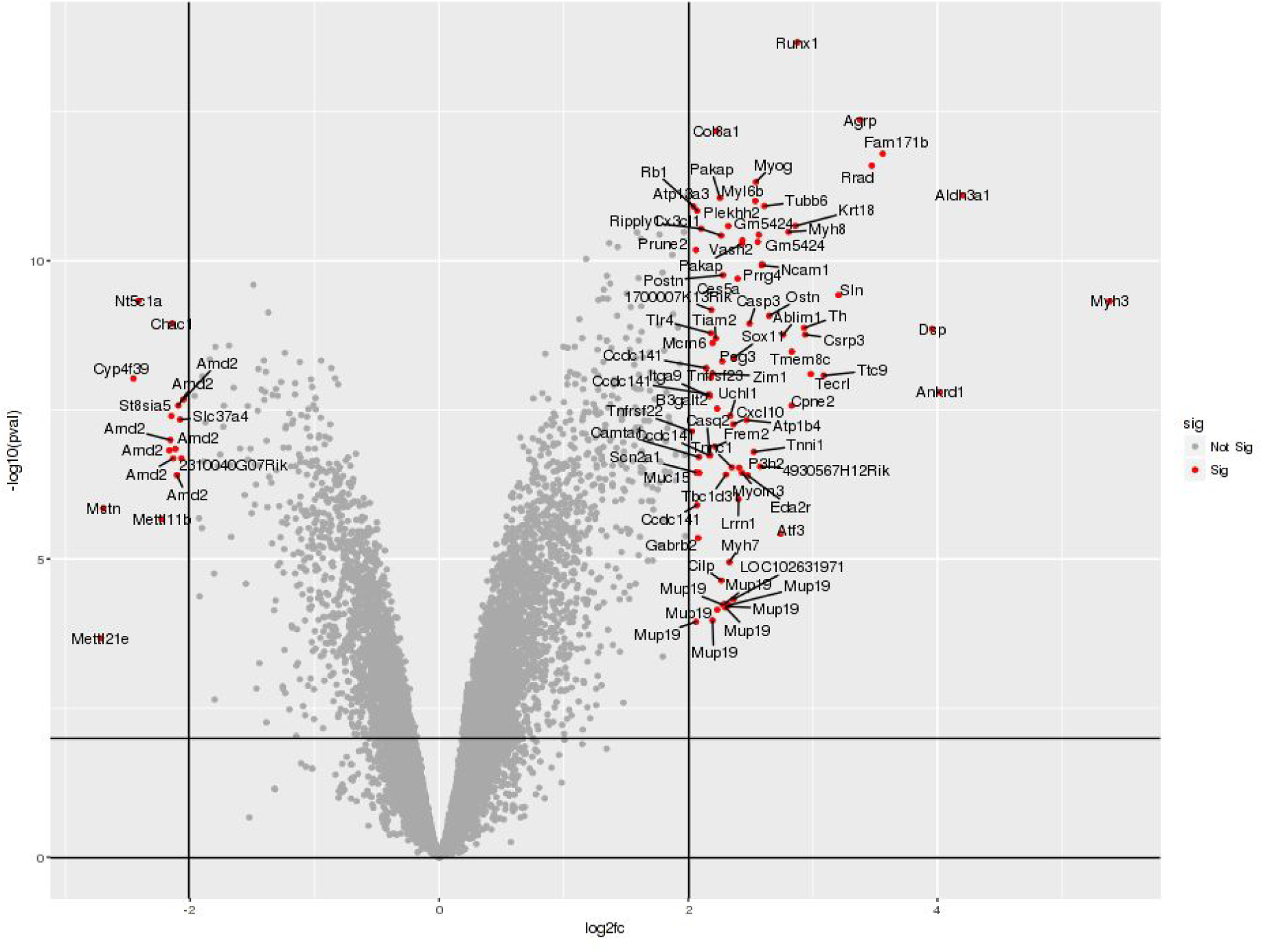

| ID | Description | GeneRatio | BgRatio | pvalue | p.adjust |
| --- | --- | --- | --- | --- | --- |
| GO:0002318 | myeloid progenitor cell differentiation | 5/124 | 16/30953 | 4.010838e-09 | 4.024227e-06 |
| GO:0006941 | striated muscle contraction | 12/124 | 315/30953 | 5.769501e-09 | 4.024227e-06 |
| GO:0070252 | actin-mediated cell contraction | 9/124 | 166/30953 | 2.471387e-08 | 1.149195e-05 |
| GO:0048738 | cardiac muscle tissue development | 13/124 | 452/30953 | 3.652707e-08 | 1.170199e-05 |
| GO:0043501 | skeletal muscle adaptation | 6/124 | 49/30953 | 4.443638e-08 | 1.170199e-05 |

KEGG enrichment analysis Show entries
| ID | Description | GeneRatio | BgRatio | pvalue | p.adjust |
| --- | --- | --- | --- | --- | --- |
| mmu04260 | Cardiac muscle contraction | 9/90 | 124/10680 | 9.170735e-07 | 0.0001430635 |
| mmu04261 | Adrenergic signaling in cardiomyocytes | 10/90 | 227/10680 | 1.996688e-05 | 0.0012413462 |
| mmu05410 | Hypertrophic cardiomyopathy (HCM) | 8/90 | 144/10680 | 2.743600e-05 | 0.0012413462 |
| mmu05414 | Dilated cardiomyopathy | 8/90 | 147/10680 | 3.182939e-05 | 0.0012413462 |
| mmu05145 | Toxoplasmosis | 7/90 | 175/10680 | 6.699649e-04 | 0.0179742901 |

